# Phosphoproteomic profiling highlights CDC42 and CDK2 as key players in the regulation of the TGF-β pathway in *ALMS1* and *BBS1* knockout models

**DOI:** 10.1101/2022.11.04.514246

**Authors:** Brais Bea-Mascato, Girolamo Giudice, Iguaracy Pinheiro-de-Sousa, Carlos Solarat, Evangelia Petsalaki, Diana Valverde

**Affiliations:** CINBIO, Universidad de Vigo, 36310 Vigo, Spain; Grupo de Investigación en Enfermedades Raras y Medicina Pediátrica, Instituto de Investigación Sanitaria Galicia Sur (IIS Galicia Sur), SERGAS-UVIGO, Vigo, Spain; European Molecular Biology Laboratory-European Bioinformatics Institute, Wellcome Genome Campus, Hinxton CB10 1SD, UK

**Keywords:** ALMS1, BBS1, ciliopathies, TGF-β, phosphoproteomics, Alström syndrome, Bardet-Biedl syndrome.

## Abstract

**BACKGROUND:** The primary cilium is a sensory organelle that extends from the plasma membrane. It plays a vital role in physiological and developmental processes by controlling different signalling pathways such as WNT, Sonic hedgehog (SHh), and transforming growth factor β (TGF-β). Ciliary dysfunction has been related to different pathologies such as Alström (ALMS) or Bardet-Biedl (BBS) syndrome. The leading cause of death in adults with these syndromes is chronic kidney disease (CKD), which is characterised by fibrotic and inflammatory processes often involving the TGF-β pathway.

**METHODS:** Using genomic editing with CRISPR-CAS9 and phosphoproteomics we have studied the TGF-β signalling pathway in knockout (KO) models for *ALMS1* and *BBS1* genes. We have developed a network diffusion-based analysis pipeline to expand the data initially obtained and to be able to determine which processes were deregulated in TGF-β pathway. Finally, we have analysed protein-protein and kinase-substrate interactions to prioritise candidate genes in the regulation of the TGF-β pathway in ALMS and BBS.

**RESULTS:** Analysis of differentially phosphorylated proteins identified 10 candidate proteins in the *ALMS1* KO model and 41 in the *BBS1* KO model. After network expansion using a random walk with a restart algorithm, we were able to identify the TGF-β signalling pathway together with other related processes such as endocytosis in the case of *ALMS1* or the regulation of the extracellular matrix in *BBS1*. Protein interaction analyses demonstrated the involvement of CDC42 as a central protein in the interactome in *ALMS1* and CDK2 in the case of *BBS1*.

**CONCLUSION:** In conclusion, the depletion of *ALMS1* and *BBS1* affects the TGF-β signalling pathway, conditioning the phosphorylation and activation of several proteins, including CDC42 in the case of *ALMS1* and CDK2 in the case of *BBS1*.

## INTRODUCTION

The primary cilium is a microtubule-based sensory organelle that extends from the plasma membrane. It is present in all cell types and plays a vital role in physiological and developmental processes by controlling different signalling pathways such as WNT, Sonic hedgehog (SHh), and transforming growth factor β (TGF-β) (Berbari et al., 2009; Ishikawa and Marshall, 2011; Christensen et al., 2012; May-Simera et al., 2018; Anvarian et al., 2019). Ciliary dysfunction generates different phenotypes, the most common of which are obesity, type 2 diabetes mellitus, retinopathies or the appearance of multi-organ fibrosis leading to liver, kidney, or lung dysfunction (Gerdes et al., 2009; Waters and Beales, 2011; Reiter and Leroux, 2017; May-Simera et al., 2018).

The group of diseases characterised by ciliary dysfunction, mutations in primary cilium or basal body genes, are commonly known as ciliopathies. Alström syndrome (ALMS) and Bardet-Biedl syndrome (BBS) are two model ciliopathies where, despite the progress made in recent years, a clear pathological mechanism has not yet been established. (Forsythe and Beales, 2012; Forsythe et al., 2018; Hearn, 2018). ALMS is a monogenic recessive disease with an incidence of 1 in 1,000,000 individuals in the general population. To date, the only known causative gene is the *ALMS1* gene, which codes for a centrosome-associated protein, responsible for maintaining cohesion between the mother centriole and the daughter centriole (Collin et al., 2002; Hearn et al., 2005; Marshall et al., 2011). On the other hand, BBS is also an autosomal recessive disorder, with 23 causative genes described to date, named from *BBS1* to *BBS23*, being BBS1 the most prevalent gene with pathogenic variants in 23% of described cases (Forsythe and Beales, 2012; Forsythe et al., 2018; Perea-Romero et al., 2022). The main proteins of BBS, BBS1, BBS2, BBS4, BBS5, BBS7, BBS8, BBS9 and BBS18/BBSIP1, encode for an octameric protein complex called the BBSome involved in primary cilia transport (IFT), retrograde and anterograde (Klink et al., 2020; Singh et al., 2020). The clinical phenotypes of these two ciliopathies are overlapping, making their differentiation difficult and complicating the correct diagnosis (Aliferis et al., 2012). One of the main causes of death in adult patients with these syndromes is chronic kidney disease (CDK) (Li et al., 2007; Marshall et al., 2011; Putoux et al., 2012; Jaykumar et al., 2018; Waldman et al., 2018; Baig et al., 2020; Meyer et al., 2022). This disease is usually accompanied by inflammatory and fibrotic processes that progressively degrade normal kidney function until eventually renal failure and death occur (Meng et al., 2014).

The TGF-β pathway is one of the main pathways involved in fibrotic and inflammatory processes at the cellular level (Lan, 2011; Finnson et al., 2020). It is related to the control of several processes such as mitosis, apoptosis, cell migration or autophagy through cross-signalling with other pathways such as MAPK, PI3K/AKT or SMADs (Massagué, 2012; Hamidi et al., 2017; Luo, 2017; Zhang et al., 2017; Gasior et al., 2019). In previous studies in our laboratory, we have determined that *ALMS1* depletion leads to altered signal transduction of the TGF-β pathway and other related pathways such as AKT or p53 (Álvarez-Satta et al., 2021; Bea-Mascato et al., 2022b, 2022a), altering processes such as cell migration or EMT. In this study, we apply an original approach based on phosphoproteomics to delve into the regulation of the TGF-β pathway in ALMS and to determine for the first time whether there are alterations in this pathway in BBS.

## MATERIALS AND METHODS

### Cell culture and CRISPR knockouts

The hTERT-BJ-5ta cell line, provided by the American Type Culture Collection (ATCC), was maintained in a medium composed of 4:1 of Dulbecco’s minimal essential medium (DMEM, Gibco, Invitrogen, NY, USA) and Medium 199, Earle’s Salts (Gibco, Invitrogen, NY, USA), supplemented with 10% fetal bovine serum (FBS) (Gibco, Invitrogen, NY, USA) and 2% penicillin/streptomycin (P/S) (Gibco, Invitrogen, NY, USA). In this study, a knockout model for the *BBS1* gene was generated **(Figure S1 A, B)** using 4 different sgRNAs **(Supplementary Table S1-2)** and a knockout (KO) model for the *ALMS1* gene previously generated in Bea-Mascato et al (Bea-Mascato et al., 2022b) was used. The methodology used in the generation of the *BBS1* model was the same as described in Bea-Mascato et al (Bea-Mascato et al., 2022b).

### TGF-β stimulation and protein extraction

Cell lines were cultured in 150 mm dishes (Corning, NY, USA) until 90% confluence was reached and then serum starved for 24 hours. The following day, rhTGF-β1 ligand (R&D Systems; 240-B) was added to the plates at a final well concentration of 2ng/mL for 0 and 30 min. The plates were then washed 3 times with PBS and the cells were harvested in a volume of 1.5mL of PBS. They were then pelleted at 10000 rpm for 5 minutes in a Sigma^®^ 1-14 K at 4°C. The PBS volume was removed from the tubes and the pellets were frozen at -80C until the experiment was continued.

Protein extraction was performed by adding 500µL of TEAB 100mM lysis buffer with 1% SDS (Thermo Fisher, Waltham, USA) to each sample. Samples were sonicated by giving five 5-second pulse intervals with a wave amplitude of 20% on a Branson ultra sonicator model 102C (Branson Ultrasonic, Connecticut, USA). They were then left to stand on ice for 10 minutes with moderate agitation. Finally, the cell debris was removed by centrifuging the samples for 30 minutes at 12000rpms in a Sigma^®^ 1-14 K at 4°C. The supernatant from the tubes was transferred to low-binding tubes (Thermo Fisher, Waltham, USA) and stored at -80 degrees until the next step of the analysis. Quantification of protein lysates was performed using the Pierce™ BCA Protein Assay Kit (Thermo Fisher, Waltham, USA).

### Reduction, alkylation and precipitation

Initially, samples were centrifuged at 16,000 rpm for 10 min at 4°C. Then, 600µg per sample was transferred into a new 1.5 mL tube and adjusted to a final volume of 600µL with 100mM TEAB buffer. Samples were divided into several tubes (200µL/tube). 10µL of 200mM TCEP was added to each tube and incubated at 55°C for 1 hour. Then, 10µL of 375mM iodoacetamide (IAA) was also added to the sample and incubated for 30 minutes protected from light at room temperature (RT). Finally, 6 volumes (1200µL) of acetone pre-cooled to -20 degrees were added and left to precipitate overnight. The next day the samples were centrifuged at 8000rpms for 12 min at 4°C, the acetone was decanted off and the pellet was left to dry for 2-3 min.

### Protein digestion and cleaning, TiO_2_ enrichment, TMT labelling and LC-MS/MS analysis

Approximately 200µg of protein pellets precipitated in acetone were resuspended with 200 µL of 100mM TEAB buffer. Then 5µL (5µg) of trypsin was added per tube and digested at 37°C overnight. The next day, 10 µL of 5% trifluoroacetic acid (TFA) was added to reduce the pH of the solution. Finally, peptide samples were cleaned up using C18 desalting tips (Agilent OMIX 100 µL, SPE with pipette) and the pooled eluents with speed-vac at 45°C. Phosphopeptide enrichment was performed using TiO_2_ spin tips from the High-SelectTM TiO_2_ Phosphopeptide Enrichment Kit following the manufacturer’s protocol (Thermo Fisher, Waltham, USA). After enrichment, a further cleaning was performed using C18 desalting tips.

Samples were resuspended in 50 µL of 100mM TEAB buffer and 20 µL of TMT label reagent was added to each sample, using TMT10plex™ Isobaric Mass Tagging Kit (Thermo Fisher, Waltham, USA). Samples were multiplexed and analyzed by LC-MS/MS using a 90-minute gradient on Orbitrap Eclipse (Thermo Fisher, Waltham, USA) with a method using CID-MSA for the identification of MS2 and a real-time search algorithm to quantify reporters in MS3. As a quality control, the BSA controls were digested in parallel and ran between each of their samples to avoid carryover and assess instrument performance.

### Protein identification and differential phosphorylation analysis

The identification of peptides/proteins was performed using the software Proteome Discoverer software (v2.5) **(Supplementary Table S3-4)**. Samples were searched against the SP_Human database (March 2021), using the search algorithm Mascot v2.6 (http://www.matrixscience.com/). Peptides have been filtered based on false discovery rate (FDR) and only peptides showing an FDR lower than 5% have been retained. The peptides were grouped into proteins with the default option of Proteome discovery (v2.5) and these were used in the downstream analysis.

The proteins resulting from the identification were used for differential phosphorylation analysis. Each multiplexed/batch (with and without stimulation of the TGF-β pathway) was analysed individually with the R (v4.2.1) package DEP (Zhang et al., 2018). The data was filtered to keep only the proteins that were quantified in all samples of at least one condition. Subsequently, the normalization of the data by variance stabilizing normalization (VSN) was performed. Finally, for data imputation, the MiniProb algorithm was used with a q-value of 0.01. The log2 fold change (FC) of the differentially phosphorylated proteins was calculated against the controls (WT) of the corresponding batch for both knockouts (*ALMS1* and *BBS1*) **(Supplementary Table S5-6)**. For downstream analyses, the differentially phosphorylated protein was that with a value of |log2FC| > 0.5 and an FDR < 0.05.

### Network diffusion, enrichment analysis and protein-protein interaction analysis

For network diffusion analysis, a graph covering the entire human signalome was created with the R package, igraph (Csardi and Nepusz, 2006). The creation of the human interactome was performed as follows. Initially, the human interactome was extracted from the IntAct database (version: 4.2.17, last updated May 2021) (Orchard et al., 2014). In addition, kinase-substrate interactions and kinase-kinase interactions contained in PhosphoSitePlus (Orchard et al., 2014) (version 6.5.9.3, last update May 2021), OmniPath (Türei et al., 2016) (last version May 2021) and SIGNOR 2.0 (Licata et al., 2020) (last version May 2021) were integrated in this interactome. All proteins not annotated in Swiss-Prot (UniProt, 2021) and those not annotated with at least one GO term (Gene Ontology, 2021) were removed. The resulting protein interaction network (PIN) comprised a total of 16,407 nodes and 238,035 edges. The weight of edges was defined by the semantic similarity between gene pairs, modelled according to Topological Clustering Semantic Similarity (TCSS) algorithm (Jain and Bader, 2010) and calculated using the Semantic Measure Library (Harispe et al., 2014). The raw PIN was normalised to avoid hub bias using “correct.for.hubs” option in diffusR.

The differentially phosphorylated proteins detected in the “*Protein identification and differential phosphorylation analysis*” section, were used as seed nodes in the interactome graph. These nodes were assigned the same weight and a weight of 0 to the rest of the network. Finally, the random walk with restart (RWR) algorithm (Tong et al., 2006; Köhler et al., 2008) was used through the R package diffusR using a restart probability of 0.3, 10,000 iterations and a threshold of 1e-06. Nodes were sorted by RWR score after diffusion, and to ensure at least 10% real (not inferred) data, the top 100 were used in subsequent analyses **(Supplementary Table S7-8)**.

The enrichment analysis was performed using the R package enrichR, an R interface for accessing the enrichR database (Kuleshov et al., 2016). The differentially phosphorylated proteins and the 100 nodes with the highest RWR scores were compared with the GO Molecular Function 2021, GO Biological Process 2021 and BioPlanet 2019 datasets in independent enrichment analysis. Only the 20 processes with the lowest FDR were considered using an FDR < 0.05 as a threshold.

The protein-protein interaction results obtained after network diffusion for the top 100 proteins was visualised with the help of Cytoscape software (v 3.9.1) (Shannon et al., 2003).

Lastly, a kinase-substrate enrichment analysis (KSEA) was performed using the KEA3 web application (Kuleshov et al., 2021). The 100 proteins with the highest RWR score were used as input. Both rows and columns of the graph were clustered using rank by sum to obtain the top substrate and top kinase in the first row and column of the heatmap.

## RESULTS

### *ALMS1* and *BBS1* depletion alters the phosphorylation of several proteins upon TGF-β1 ligand stimulation

To identify whether TGF-beta signalling was affected in the ALMS1 and BBS1 knockout cells and discover potentially new processes, we performed global phosphoproteomics in the presence and absence of TGF-β1 stimulation **(methods)**. The experiment was designed using two multiplexes of 10 samples each. The batch 1 contains all samples without TGF-β1 stimulation, while batch 2 contains all samples stimulated for 30 minutes with TGF-β1 **(Figure 1A-B)**. We identified a total of 2,720 phosphopeptides (1,406 proteins) in batch 1 (samples without stimulation) and 3,317 phosphopeptides (1,619 proteins) in batch 2 (samples with stimulation), 1076 common proteins were identified in both batches **(Figure 1B)**. Of the total peptides/proteins identified, 50-60% were quantified in all samples of at least one condition (731 proteins in batch 1 and 960 in batch 2) **(Figure S1 C-F)**.

**Figure 1.**
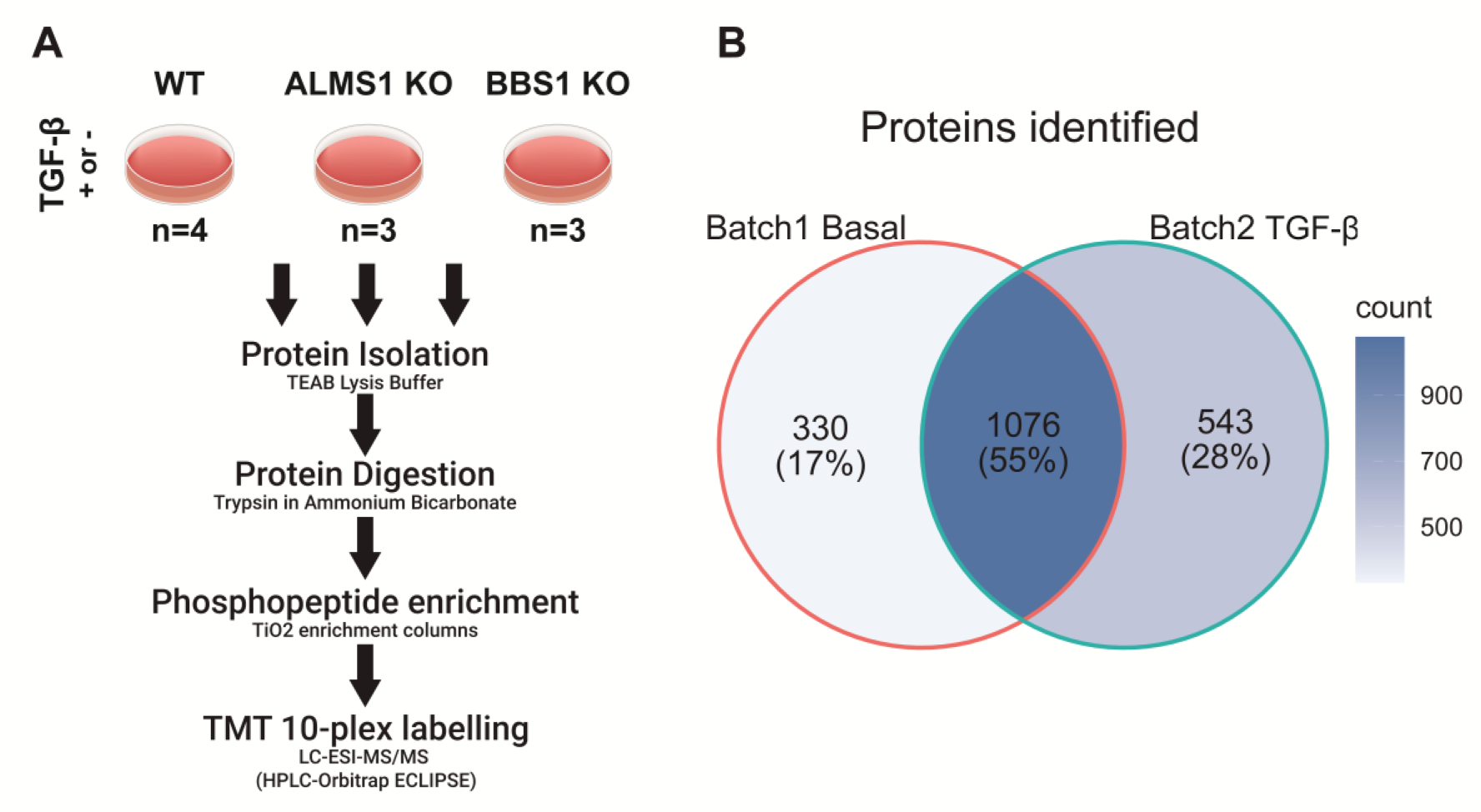
Experimental design of the phosphoproteome in basal and TGF-β-stimulated BJ-5ta. **(A)** Protocol followed for the generation of the phosphoproteomic datasets **(B)** Overlap between the proteins identified in Batch 1 (Basal) and Batch 2 (TGF-β).

After differential phosphorylation analysis, no significant proteins were detected in batch 1 for *BBS1* knockout (BBS1KO) and 3 differentially phosphorylated proteins (DPP) were detected in the case of *ALMS1* knockout (ALMS1KO) **(Figure 2A-C)**. Results from batch 2 showed 41 DPP in BBS1KO and 10 in ALMS1KO **(Figure 2B-D)**. Of the total DPPs detected in the cell models, 4 were common among them, 3 of them were infra-phosphorylated (EHBP1, CDC42EP3, RPL27A) while 1 (SLC38A1) was over-phosphorylated.

**Figure 2.**
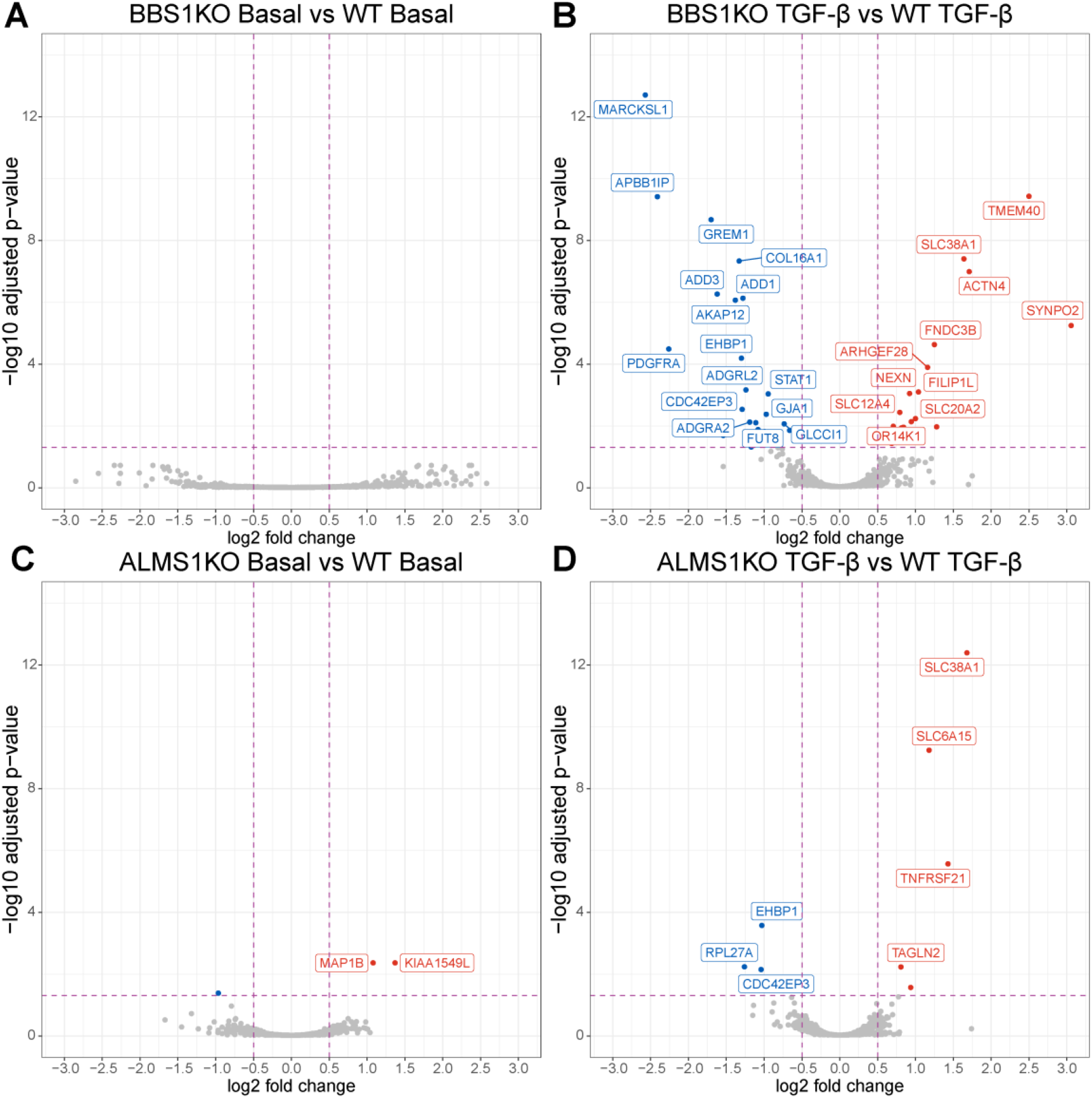
Differential phosphorylation analysis showed alterations in protein activation after TGF-β stimulation. Labelled proteins have an FDR < 0.01. **(A)** Volcano-plot of differentially phosphorylated proteins (DPP; FDR < 0.05) in *BBS1* knockout before TGF-β stimulation **(B)** Volcano-plot of DPP in *BBS1* knockout after TGF-β stimulation **(C)** Volcano-plot of DPP in *ALMS1* knockout before TGF-β stimulation **(D)** Volcano-plot of DPP in *ALMS1* knockout after TGF-β stimulation.

### Network diffusion highlights the involvement of *ALMS1* and *BBS1* in processes coordinated by the TGF-β pathway

Then, we performed a pathway enrichment analysis **(methods)** to identify the processes that are most affected by *ALMS1* and *BBS1* depletion after TGF-β stimulation **(Figure 3)**. Pre-diffusion ORA was performed using the DPP after TGF-β stimulation (only seed nodes). Due to the small number of proteins and the low signal-to-noise ratio of phosphoproteomics, we were unable to identify the TGF-β pathway, which serves as a control of the analysis **(Figure 3A, C)**. To improve the signal-to-noise ration from our DPP, we applied a network diffusion-based analysis **(methods)**. The ORA after diffusion was performed with the top 100 proteins with the highest RWR score **(Figure 3B, D)**. In this case, we obtained TGF-β in both KO cell models. In addition, processes related to TGF-β regulation such as mitosis, NOTHC1, p53 or SHh signalling in the case of *BBS1KO* and endocytosis or Syndecan4 in the case of *ALMS1KO* were also identified **(Figure 3B, D)**.

**Figure 3.**
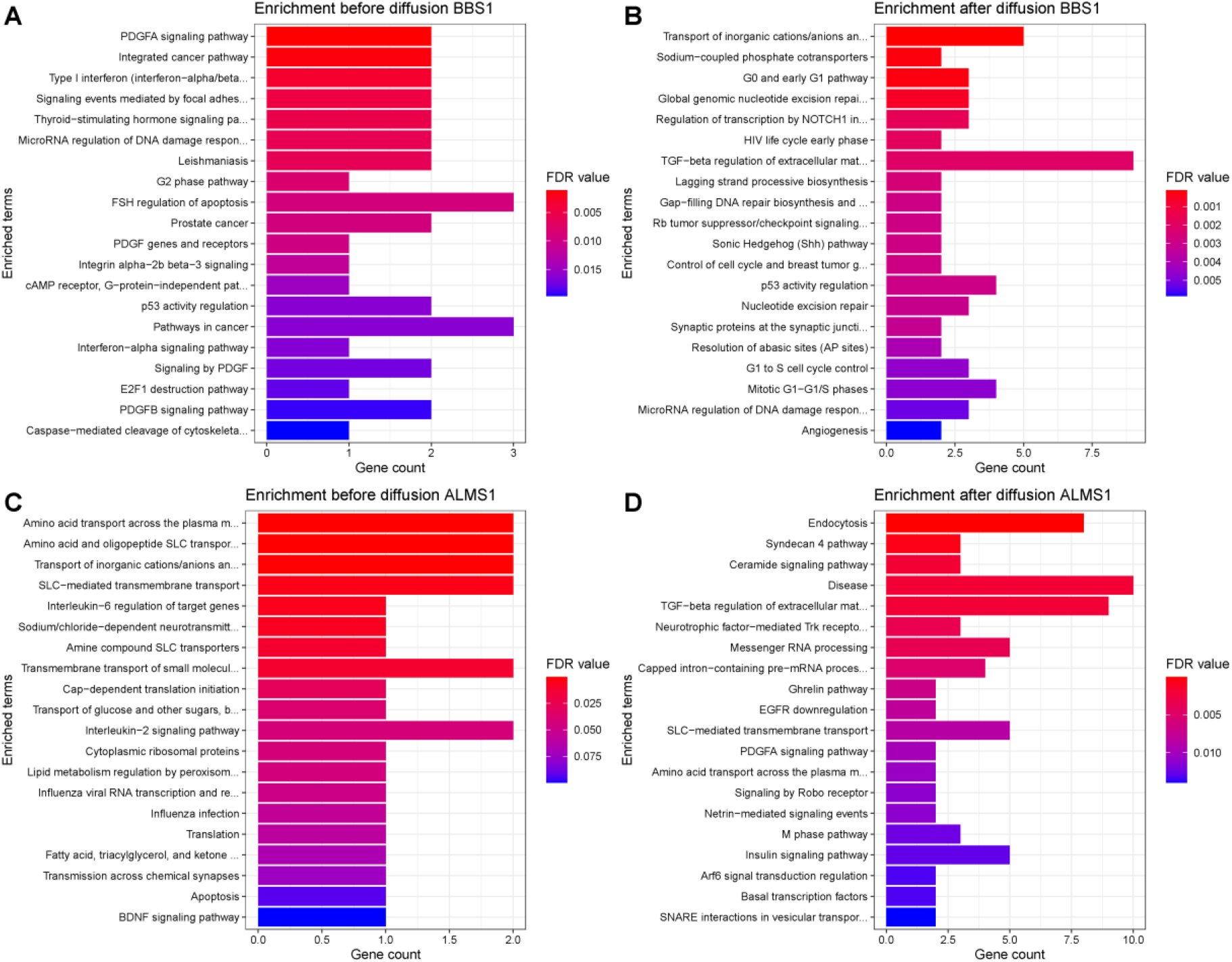
Over-representation analysis (ORA) after network diffusion reveals processes controlled by TGF-β such as endocytosis, mitosis, or extracellular matrix in Bioplanet DB. **(A)** ORA of DPP in the *BBS1*KO before network diffusion. **(B)** ORA of top 100 proteins with the highest RWR score in the *BBS1*KO after network diffusion. **(C)** ORA of DPP in the *ALMS1*KO before network diffusion. **(D)** ORA of top 100 proteins with the highest RWR score in the *ALMS1*KO after network diffusion.

### Analysis of protein-protein interactions involves CDC42 and several RAB proteins in the alterations of TGF-β signalling in *ALMS1* and CDK2 in *BBS1*

Using the top 100 proteins with the highest RWR score, an analysis of protein-protein interactions was performed. In the case of *BBS1KO*, CDK2 (seed node), a key regulator of the cell cycle, was one of the most interconnected nodes, highlighting its central role in this interactome **(Figure 4A)**. On the other hand, in the case of *ALMS1KO*, CDC42EP3 a direct effector of CDC42, key regulator of cell cycle and migration processes, was found to be one of the most interconnected nodes in the network, highlighting its central role in the deregulated interactome of this model. It also highlights the highly interconnected cluster of RAB proteins such as RAB11 and RAB8A, resulted by EHBP1, that play an important role in endocytosis **(Figure 4B)**.

**Figure 4.**
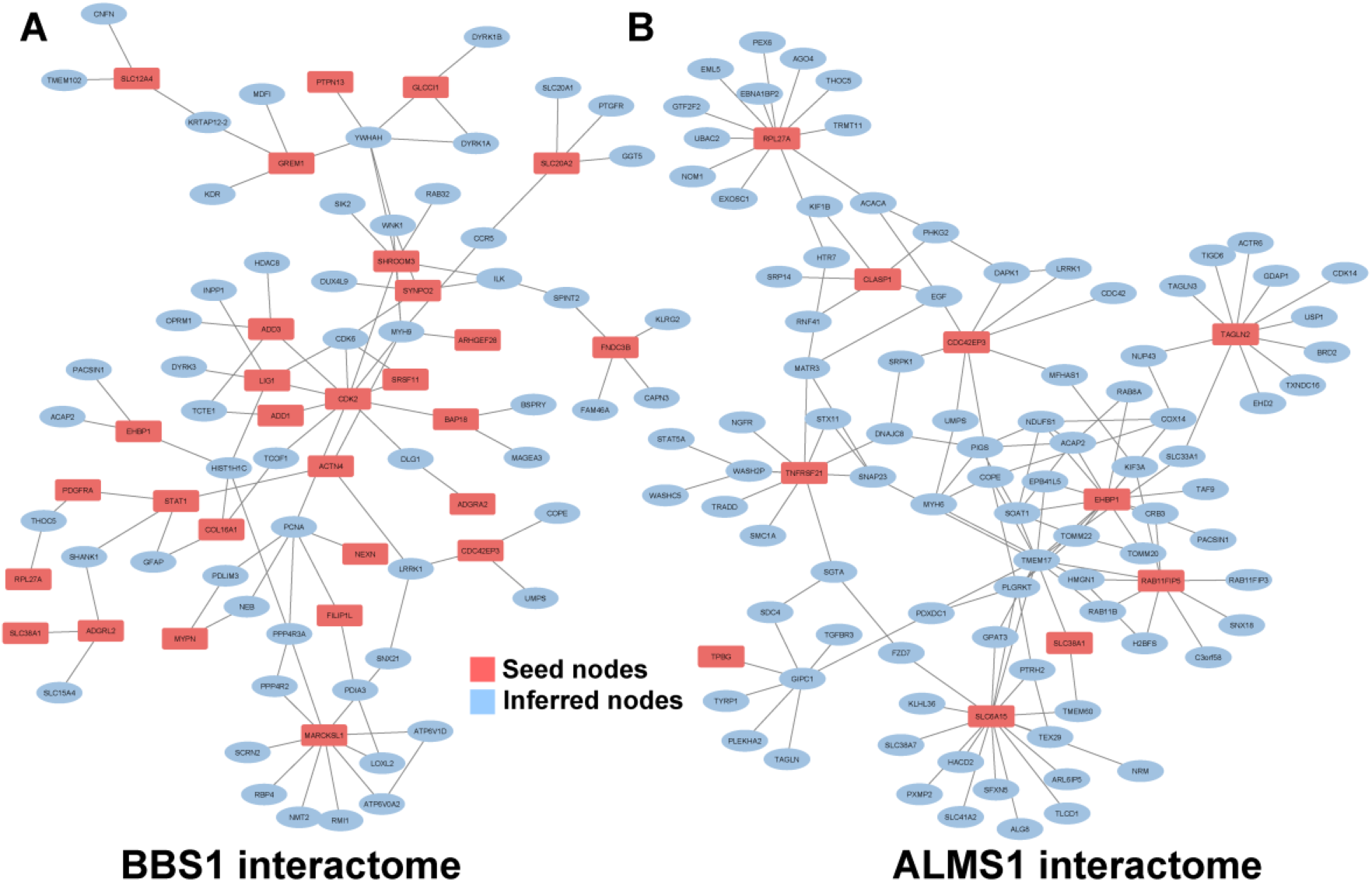
Protein-protein interaction analysis highlights CDK2 as the central node of the *BBS1* interactome and CDC42EP3 as the central node of the *ALMS1* interactome. Red nodes represent the DPP that were detected in the first analysis. Blue nodes represent the nodes inferred by RWR algorithm. **(A)** Interactome of the top 100 proteins with the highest RWR score after network diffusion in the *BBS1KO*. **(B)** Interactome of the top 100 proteins with the highest RWR score after network diffusion in the *ALMS1KO*.

### Kinase-substrate enrichment analysis uncovers CDK2 and CDC42 as the most phosphorylated substrates

To determine which DPP were subject to the highest level of up-regulation, a KSEA was performed with the top 100 with the highest RWR score. The proteins (substrates) subject to the highest level of up-regulation were again CDK2 in the case of *BBS1* and CDC42 in the case of *ALMS1* **(Figure 5A-B)**. The main kinases regulating the selected subset of proteins were AKT1 in *BBS1* and CDK2 in the case of *ALMS1* **(Figure 5A-B)**. This shows that the observed alterations have commonalities but appear to differ in their regulation.

**Figure 5.**
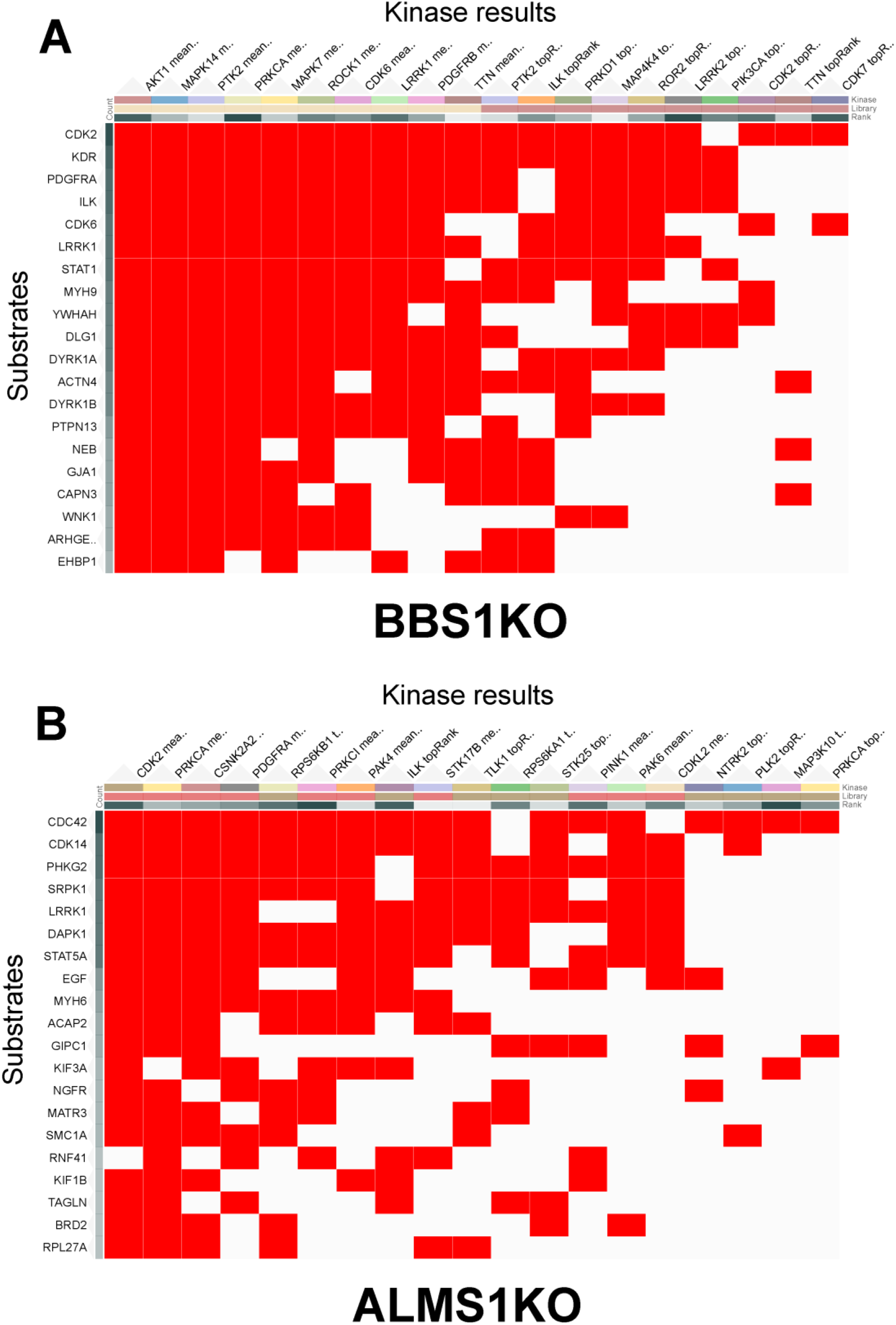
KSEA highlights CDK2 in *BBS1* and CDC42 in *ALMS1* as the main substrates subject to regulation. **(A)** KSEA of the top 100 proteins (substrates) with the highest RWR score after network diffusion in the *BBS1KO*. **(B)** KSEA of the top 100 proteins (substrates) with the highest RWR score after network diffusion in the *ALMS1KO*. The plots only show the 20 most up-regulated proteins (substrates) out of the initial 100 proteins used in the analysis.

## DISCUSSION

The TGF-β pathway is one of the main signalling pathways regulated by the primary cilium (Clement et al., 2013; Pedersen et al., 2016; Christensen et al., 2017; Anvarian et al., 2019). It controls processes related to cell proliferation and migration and is associated with several pathologies such as liver or kidney fibrosis, recurrent phenotypes in ciliopathies (Zhang et al., 2017; Guan et al., 2021; McConnachie et al., 2021). Despite this, few studies have attempted to link ciliary gene dysfunction with signalling alterations in the TGF-β pathway (Mönnich et al., 2018; Álvarez-Satta et al., 2021; Bea-Mascato et al., 2022b, 2022a). In this study, we have applied phosphoproteomics to determine alterations in the TGF-β signalling pathway in two model ciliopathies, ALMS and BBS. We have further elucidated the regulation of the TGF-β pathway in ALMS and established for the first time the link between the TGF-β pathway and BBS genes (Álvarez-Satta et al., 2021; Bea-Mascato et al., 2022b, 2022a).

In this article, we designed and applied a pipeline that allowed us to obtain a global view of the processes altered in the TGF-β pathway by inhibiting two ciliary genes, *ALMS1* and *BBS1*. In the case of *ALMS1*, we found that the altered processes were mainly related to endocytosis and mitosis (*via* the Syndecan 4 signalling pathway, responsible for cytokinesis) (Keller-Pinter et al., 2010). The role of *ALMS1* in endocytotic and mitotic processes has already been established in the past (Zulato et al., 2011; Collin et al., 2012; Leitch et al., 2014; Jaykumar et al., 2018; Bea-Mascato et al., 2022b). However, this is the first time that these alterations in endocytosis and mitosis could be related to the TGF-β signalling pathway affecting the activation of RAB11FIP5 and EHBP1 (seed nodes) and other related proteins such as RAB8A and RAB11B (inferred nodes) among others (Rai et al., 2020; Solinger et al., 2020). In addition, the relationship of CDC42 in this interactome has also been established through CDC42EP3, a CDC42 effector protein that is under-phosphorylated in the differentially phosphorylated proteins detected in the differential analysis **(Figures 4B, 2D)**. The significance of CDC42 has also been determined by KSEA **(Figure 5B)**. CDC42 plays a key role in cell polarity and regulation of mitosis (Witte et al., 2017; Moran et al., 2019; Rich-Robinson et al., 2021). CDC42EP3 acts downstream of CDC42 and regulates the rearrangement of the actin and septin cytoskeleton (Farrugia and Calvo, 2016). Cell cycle disorders and cytoskeleton alterations are common phenotypes in *ALMS1*-depleted cells, so a dysfunction of CDC42 and CDC42EP3 could be a possible cause of them (Zulato et al., 2011; Collin et al., 2012; Shenje et al., 2014; Bea-Mascato et al., 2022b). Inhibition of CDC42EP3 and EHBP1 in *BBS1*KO could indicate commonalities in regulation between *ALMS1* and *BBS1*, so that the alterations detailed above could also be present when BBS1 is inhibited.

*BBS1* gene inhibition altered the phosphorylation of 41 proteins after TGF-β stimulation. Following network diffusion, these proteins were found to be mainly involved in the regulation of the extracellular matrix *via* the TGF-β pathway; cross-signalling with other pathways such as p53, NOTCH1 or SHh and transport of cation ions (Ca^2+^ or Na^+^) **(Figure 3B)**. Alterations of TGF-β signalling have already been described when inhibiting other ciliary genes such as *ALMS1* or *CEP128*, but never for the *BBS1* gene (Mönnich et al., 2018; Álvarez-Satta et al., 2021; Bea-Mascato et al., 2022b). Some of the main nodes of the BBS1 interactome and KSEA were SYNPO2, CDK2 and PDGFRA proteins. SYNPO2, which is over-phosphorylated, encodes actin binding proteins that has been characterized as a tumor suppressor (OuYang et al., 2021). It is considered a protein with inhibitory capacity on cell migration and proliferation (OuYang et al., 2021). On the other hand, CDK2 and PDGFRA, which appear under-phosphorylated, are key regulators of mitosis and cell proliferation (Elling et al., 2011; Spencer et al., 2013; Li et al., 2018). Thus, the lack of BBS1 seems to prevent proper TGF-B-mediated proliferation. Our data do not allow us to conclude the existence of a compensatory mechanism via another pathway.

The main limitations of this study are related to the use of in vitro models, which often do not fully reproduce patient phenotypes. For this reason, these results need to be validated in primary samples or animal models to determine whether the alterations described are conserved processes.

On the other hand, phosphoproteomics is an approach that allows the global analysis of the activation of the cellular interactome. However, like any technique, it has several limitations, since the phosphoproteome is estimated at 30% of the total proteome and current enrichment techniques have a recovery of 1-3% of the total amount of starting sample (Low et al., 2021; Urban, 2022). The first limitation is due to a large amount of starting material needed to obtain the first data. The recommended starting material required is usually a minimum of 600µg-1mg of protein lysate (Solari et al., 2015). This is affordable when working with cell culture, but very complicated when working with tissue samples. This makes it necessary to develop techniques that increase the enrichment yield and allow the use of less starting material. Phosphoproteomics data are usually scarce and noisy, requiring the development of statistical techniques and analysis methodologies that allow robust inference from the data initially obtained (Solari et al., 2015). Furthermore, the nature of the data generated by current phosphoproteomics techniques introduces a bias in that kinases have only been identified from less than 5% of the phosphoproteome, and functional assignments of phosphosites are almost negligible (Needham et al., 2019).

## CONCLUSION

In conclusion, the depletion of ciliary genes such as *ALMS1* and *BBS1* alters signal transduction through the TGF-β pathway, altering processes such as endocytosis and mitosis in the case of *ALMS1* and the regulation of the extracellular matrix, p53, NOTCH1, SHh and transport of cation ions in the case of *BBS1*. These phenomena could be explained by alterations affecting the inhibition of CDCD42 in the case of *ALMS1* or CDK2 in the case of *BBS1*.

## Author Contributions

BB-M, DV designed the study. BB-M performed the experiments and analysis. EP led the design of the analysis pipeline for differential phosphorylation and network diffusion. GG and IP-S supervised and assisted in the analysis. CS performed the screening and characterisation of clones for the *BBS1* KO model. All authors wrote the manuscript and provided approval for publication.

## Informed Consent Statement

Not Apply.

## Data Availability Statement

Data are available via ProteomeXchange with the identifier PXD037573.

## Code Availability Statement

The code used in this study can be consulted on the GitHub repository https://github.com/BreisOne/phosphoproteome_pipeline.

## Acknowledgements

We sincerely thank the Proteomics and Genomics services from Centro de Apoyo Científico-Tecnológico a la Investigación (CACTI) of University of Vigo and its specialists Paula Álvarez Chaver, Ángel Sebastián Comesaña, Verónica Outeiriño and Manuel Marcos for their guidance and advise. We also thank Mercedes Peleteiro Olmedo from Centro de Investigacións Biomédicas (CINBIO) from University of Vigo for the flow cytometry service. Finally, thanks to Eduard Sabido and Cristina Chiva Rodríguez from the Centre for Genomic Regulation (CRG) of the Pompeu Fabra University (UPF) for the data acquisition service and their advice and guidance in optimising the process.

