## Supplementary material for "Phosphoproteomic profiling highlights CDC42 and CDK2 as key players in the regulation of the TGF-β pathway in *ALMS1* and *BBS1* knockout models"

CINBIO Facultad de Biología, Universidad de Vigo, Campus As Lagoas-Marcosende s/n, 36310 Vigo, Spain

### Supplementary Material:

**Supplementary Table S1.** sgRNAs used to generate the KO model for the BBS1 gene. The letters in red signify the base pairs added for ligation of the oligos in the LentiCRISPRv2 plasmid. The sequence of the sgRNA is on the forward strand. The reverse strand sequence is shown for the generation of the double-stranded oligo to be inserted into the plasmid. In addition, the sgRNAs are shown with the off-target effect score and the on-target effect score.

| | ID | Sequence | On-target Score | Off-target Score | PAM | Strand |
|---|---|---|---|---|---|---|
| gRNA 1 | gRNA 1.BBS1Exon 9. FWD | CACCGTGTTGAGTTCCGGCTTGCCG | 61,7 | 92 | CGG | + |
| | gRNA 1.BBS1Exon 9. RVS | aaacCGGCAAGCCGGAACTCAACAC | | | | |
| gRNA 2 | gRNA 2.BBS1Exon 1. FWD | CACCGCAGGCGTCGGAATCCGATG | 69,2 | 92,4 | AGG | - |
| | gRNA 2.BBS1Exon 1. RVS | aaacCATCGGATTCCGACGCCTGC | | | | |
| gRNA 3 | gRNA 3.BBS1Exon 10. FWD | CACCGTGTACCCGGATAAGTCCCAC | 65 | 83 | AGG | - |
| | gRNA 3.BBS1Exon 10. RVS | aaacGTGGGACTTATCCGGGTACAC | | | | |
| gRNA 4 | gRNA 4.BBS1Exon 4. FWD | CACCGGGCTTTCGGTCATCACCAG | 71,3 | 75,3 | TGG | - |
| | gRNA 4.BBS1Exon 4. RVS | aaacCTGGTGATGACCGAAAGCCC | | | | |

**Supplementary Table S2.** Primers used for the characterisation of the mutations generated by the sgRNAs in the BBS1 gene. TM calculated with TM calculator (Thermo Fisher, Waltham, USA) and TM optimised empirically.

| BBS1 | Forward | Reverse | Amplified fragment (pb) | Tm calculated | Tm optimised |
|---|---|---|---|---|---|
| Exon 1 | TGAGCCTGGGTGGGAAAG | GCCTCGGTTTCCCTATCT | 297 | 59.8 | 60.5 |
| Exon 4 | AGGTCAGGCAGCTTGAAA | AGTGCCAGAGCTGGGGTC | 300 | 58.5 | 63 |
| Exon 9 | TGGGACTTTAGACCAGGCAC | AAAGCCCACTCTCATCTTGC | 293 | 56.9 | 59 |
| Exon 10 | AAGGTGGCAGAAGTGGAAAT | AGGGAACAGCCAGCGTCT | 414 | 56.2 | 60.5 |

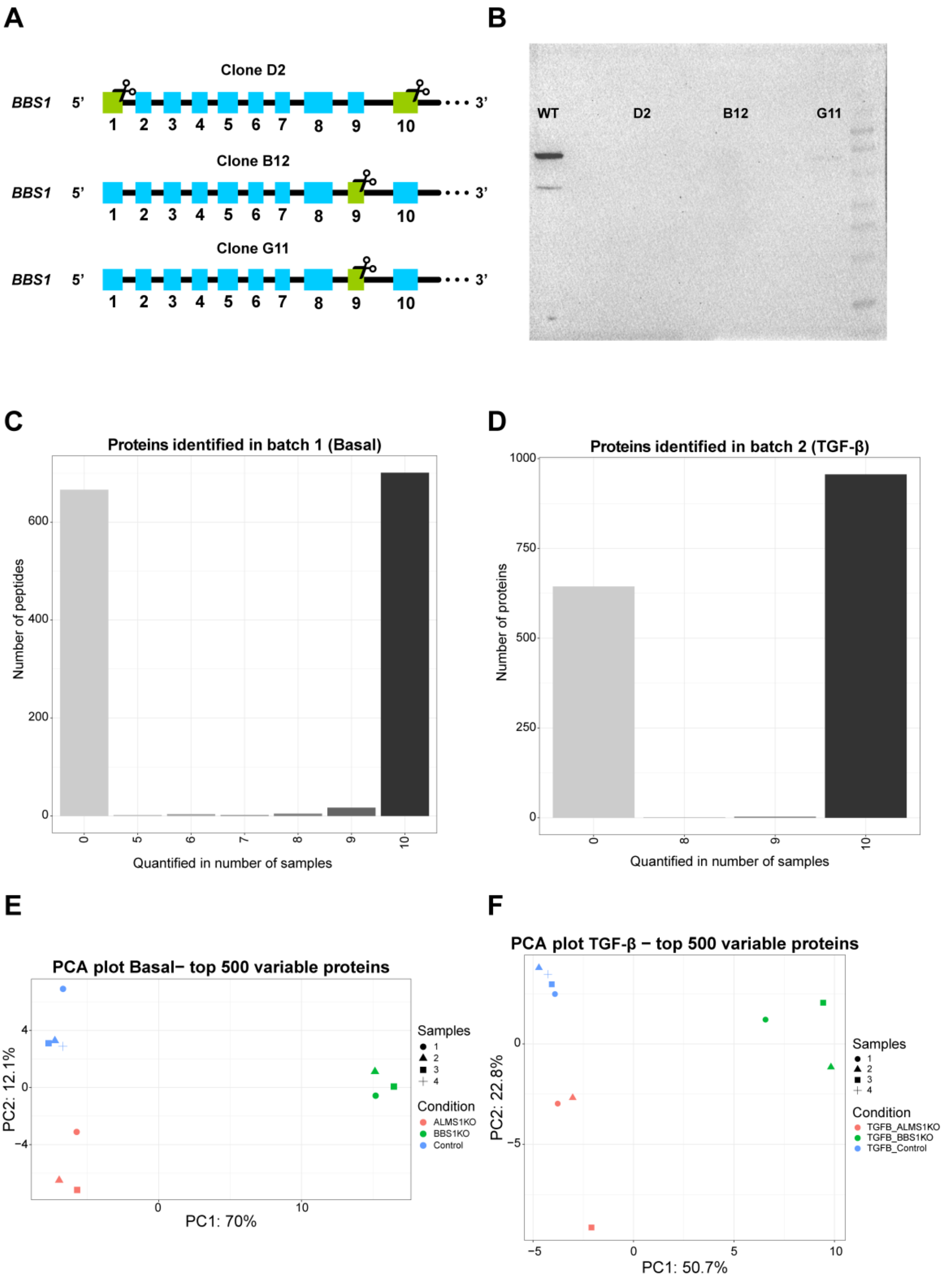
A
B
Clone D2
BBS1 5'
1 2 3 4 5 6 7 8 9 10
3'
Clone B12
Clone G11
WT
D2
B12
G11
C
Proteins identified in batch 1 (Basal)
Number of peptides
0
200
400
600
0 5 6 7 8 9 10
Quantified in number of samples
D
Proteins identified in batch 2 (TGF-β)
Number of proteins
0
250
500
750
1000
0 8 9 10
Quantified in number of samples
E
PCA plot Basal− top 500 variable proteins
PC2: 12.1%
PC1: 70%
Samples
1
2
3
4
Condition
ALMS1KO
BBS1KO
Control
F
PCA plot TGF-β − top 500 variable proteins
PC2: 22.8%
PC1: 50.7%
Samples
Condition
TGFB_ALMS1KO
TGFB_BBS1KO
TGFB_Control

**Supplementary Figure S1.** Characterisation of the *BBS1* knockout model and quality control of phosphoproteomic datasets **(A)** Graphical representation of the mutated exon/s in the 3 knockout clones obtained after screening for the *BBS1* model. **(B)** Western-blot validation of the BBS1 knockout clones (D2, B12, G11) **(C)** Number of proteins quantified in the different samples composing batch 1 (Basal) out of the total number of proteins identified **(D)** Number of proteins quantified in the different samples composing batch 2 (TGF-β) out of the total number of proteins identified **(E)** PCA plot of batch 1 (basal) with the top 500 quantified proteins with highest variation **(F)** PCA plot of batch 2 (TGF-β) with the top 500 quantified proteins with the highest variation.
